# The p97-UBXN1 complex regulates aggresome formation

**DOI:** 10.1101/2020.09.03.281766

**Authors:** Sirisha Mukkavalli, Jacob Aaron Klickstein, Betty Ortiz, Peter Juo, Malavika Raman

## Abstract

The recognition and disposal of misfolded proteins are essential for the maintenance of cellular homeostasis. Perturbations in the pathways that promote degradation of aberrant proteins contribute to a variety of protein aggregation disorders broadly termed proteinopathies. It is presently unclear how diverse disease-relevant aggregates are recognized and processed for degradation. The p97 AAA-ATPase in combination with a host of adaptor proteins functions to identify ubiquitylated proteins and target them for degradation by the ubiquitin-proteasome system or through autophagy. Mutations in p97 cause multi-system proteinopathies; however, the precise defects underlying these disorders are unclear given the large number of pathways that rely on p97 function. Here, we systematically investigate the role of p97 and its adaptors in the process of formation of aggresomes which are membrane-less structures containing ubiquitylated proteins that arise upon proteasome inhibition. We demonstrate that p97 mediates both aggresome formation and clearance in proteasome-inhibited cells. We identify a novel and specific role for the p97 adaptor UBXN1 in the process of aggresome formation. UBXN1 is recruited to ubiquitin-positive aggresomes and UBXN1 knockout cells are unable to form a single aggresome, and instead display dispersed ubiquitin aggregates. Furthermore, loss of p97-UBXN1 results in the increase in Huntingtin polyQ aggregates both in mammalian cells as well as in a *C.elegans* model of Huntington’s Disease. Together our work identifies evolutionarily conserved roles for p97 and its adaptor UBXN1 in the disposal of protein aggregates.

## Introduction

The process of protein translation, folding, and targeting to subcellular compartments is intrinsically error-prone and is therefore continuously and closely monitored by various quality control mechanisms^1–4^. Proper folding and sorting of proteins can be affected by transcription and translation errors, as well as thermal stress or pH fluctuations. As a result, an estimated 30% of newly synthesized proteins are misfolded^4,5^. Such proteins are engaged by chaperone systems to rescue functionality; however, should this approach fail, these proteins are rapidly and efficiently cleared by the ubiquitin-proteasome system (UPS) and autophagy^3^. These protein quality control mechanisms ensure that protein aggregates do not accumulate in normal cells. However aberrant accumulation of protein aggregates that are frequently ubiquitylated is a hallmark of several human disorders such as amyloidosis, diabetes, and neurodegeneration, suggesting that failure of these systems to prevent or clear the protein aggregates may have deleterious consequences^6,7^.

A variety of cellular stressors, such as proteasome impairment as examined in this study, result in a substantial increase and aggregation of misfolded, ubiquitylated proteins. These small protein aggregates are dynamically recruited to a structure known as the aggresome^8–10^. Aggresomes are membrane-less, juxta-nuclear structures encased in intermediate filament vimentin cages. In the prevailing model, ubiquitylated proteins are recognized by histone deacetylase 6 (HDAC6) and delivered by retrograde transport via the dynein motor to the microtubule organizing center^11^. Aggresomes are not mere storage sites for protein aggregates, but actively recruit chaperones, E3 ligases such as tripartite motif-containing 50 (TRIM50), the 26S proteasome, autophagy receptors such as p62 and neighbor of BRCA1 gene 1 (NBR1), and lysosomes to rescue functional, or degrade non-functional proteins^10,12,13^. Thus, aggresomes are generally believed to be cytoprotective structures that contribute to cell survival by sequestering misfolded, aggregation-prone proteins, and alleviating proteotoxic stress^14^. However, their long-term persistence is detrimental to cell viability.

Numerous studies have reported the recruitment of the p97 AAA-ATPase to aggresomes and other disease-causing protein aggregates^11,15–18^. p97 is a ubiquitin-selective unfoldase, that processes proteins for degradation by the 26S proteasome^19,20^. The functionality of p97 is best understood in its role in ER-64 associated degradation (ERAD), wherein misfolded proteins in the ER lumen are retro-translocated by p97 in an ATP-dependent manner to facilitate proteasomal degradation^21^. p97 is now appreciated to regulate extraction of ubiquitylated proteins from chromatin, ribosome-associated quality control, the cell cycle and to contribute to the selective autophagy of organelles such as mitochondria^19,22,23^. Numerous dedicated p97 adaptor proteins mediate specificity in p97 recognition of ubiquitylated substrates. The largest family of adaptors contains 13 members with conserved ubiquitin-X domains (UBXD) that bind to the N-terminus of p97^24^. Other adaptors include hetero-dimeric ubiquitin fusion degradation 1 (UFD1) and nuclear protein localization 4 (NPL4), as well as proteins that associate with the C-terminus of p97 via short linear motifs. Apart from a handful of adaptors including the UFD1-NPL4 dimer that has well-defined roles in ERAD, many adaptors remain largely unstudied and their specific roles in p97-regulated processes remain to be elucidated.

Mutations in p97 have been identified in individuals with a variety of aggregation-based disorders collectively termed multi-system proteinopathies (MSP). These include Amyotrophic Lateral Sclerosis (ALS), Inclusion Body Myopathy (IBM), Paget’s Disease (PD) and Frontotemporal Dementia (FTD), among others^25–27^. Approximately 20 of the 50 disease-associated mutations that have been identified to date span the N-terminus, linker, and first ATPase domain. This region is utilized for interaction with UBXD adaptors, and some reports suggest that these mutations impact the balance of UBXD adaptors associated with p97 and may underlie disease phenotypes^28,29^. Disease-relevant mutations in p97 impact virtually all known p97 functions, making it difficult to identify a single pathway impacted in disease. However, one striking feature common to these disorders is the accumulation and persistence of ubiquitin-positive insoluble aggregates in patient tissue.

Previous studies investigating the relationship between p97 and aggresomes reached conflicting conclusions with some studies supporting a role for p97 in aggresome formation^18,30^, while others suggesting a role in clearance^17,31^. Furthermore, the role of p97-associated adaptors has not been examined. Understanding the role of specific p97 complexes and the mechanisms by which they recognize, sequester and clear aggregates may spur the development of agents that mitigate aggregation-based disorders. We sought to address the role of p97 and the UBXD family of adaptors in the formation and subsequent clearance of aggresomes. We find that in cells with impaired proteasomal function, p97 is required for both the formation and clearance of ubiquitylated substrates at aggresomes. Importantly, we provide evidence that the p97 UBXD adaptor UBXN1 is uniquely required for the formation of aggresomes. The failure to form and clear aggresomes via p97 complexes negatively impacts cell viability. We extend these results to demonstrate that loss of the p97-UBXN1 complex results in an increase in aggregates caused by CAG (glutamine, Q) expansions (polyQ) in the Huntingtin gene both in mammalian cells as well as in a *C. elegans* model of Huntington’s Disease (HD).

Our studies have identified an evolutionarily conserved p97-dependent pathway that regulates the identification and disposal of ubiquitylated, misfolded aggregates.

## Results

### p97 is required for aggresome formation and clearance

It has been previously reported that p97 co-localizes with ubiquitin-positive aggresomes as well as aggregates caused by over-expression of disease-causing mutant proteins. However, whether p97 escorts ubiquitylated substrates to aggresomes or aids in disassembly and clearance of protein aggregates is presently unclear^17,18,30,31^. We treated cells with the proteasome inhibitor Bortezomib (Btz) in a time-course study and visualized aggregates by staining for ubiquitin. We observe aggresomes forming in the perinuclear region as early as 8 hours and by 18 hours a single large aggresome is apparent (Supplementary Figure 1A and B). We use 18 hours of Btz treatment in all following studies except in instances where we co-treated cells with the p97 inhibitor CB-5083 (Figure 1). In these experiments, aggresome formation was visualized at 8 hours as incubation with both Btz and CB-5083 for 18 hours resulted in significant cell death. Consistent with previous studies, we find that p97 is recruited to a ubiquitin-positive, perinuclear aggresome (Figure 1A). We also observe ubiquitin staining in nuclear structures resembling the nucleolus (Supplementary Figure 1A, 18 hrs). Although previous studies have reported the redistribution of ubiquitylated proteins to the nucleolus upon proteasome impairment the significance of this observation is presently unclear^32^ and nuclear aggregates were not included in our analysis. To measure cellular aggregates, their area, and spatial distribution, we developed a facile ImageJ macro called AggreCount that allows for rapid and unbiased image analysis^33^. AggreCount reports several metrics: (i) total number of cellular aggregates, (ii) percent cells with aggregates, (iii) aggregates per cell, (iv) area of aggregates, and (v) localization of aggregates (cytosol, perinuclear or nuclear). Aggresomes are identified based on (1) perinuclear localization, and (2) a minimum area cut-off that was empirically determined as the mean perinuclear aggregate area in Btz treated cells for the 8 or 18 hour time point (Supplementary Figure 1B). To investigate the role of p97 in aggresome formation we used a two-pronged approach, utilizing shRNA-mediated depletion of p97, as well as acute inhibition with CB-5083, an ATP-competitive p97 specific inhibitor. We created stable, doxycycline-inducible shRNA HeLa Flp-in TRex cell lines that allowed us to achieve transient, partial depletion of p97 without overtly impacting cell viability (Supplementary Figure 1C). Two independent sh-p97 cell lines were treated with doxycycline (to deplete p97) and Btz to promote aggresome formation, followed by release into drug-free media for a further 24 hours to monitor clearance. We imaged ubiquitin-positive aggregates by immunofluorescence and analyzed aggregates using AggreCount (Supplementary Figure 1D and E). We find that long-term p97 depletion results in an increase in the percentage of cells with aggresomes and that the aggresomes were larger (mean perinuclear aggregate area: 6.78 +/- 0.85 μm^2^), when compared to wild-type cells (mean perinuclear aggregate area: 4.2 +/- 0.15 μm^2^) (Supplementary Figure 1D and E). Notably, while wildtype cells cleared aggresomes upon removal of Btz, ubiquitin-positive aggresomes and aggregates persisted in a significant fraction of p97-depleted cells (Supplementary Figure 1D and E). In the second approach, we acutely inhibited p97 using CB-5083 concurrently with proteasome inhibition (for 8 hours) to determine the impact on aggresome formation and clearance (Figure 1B-G). Surprisingly, concurrent CB-5083 treatment with Btz resulted in a decrease in the number of cells with aggresomes compared to Btz treatment alone even though the percentage of cells with aggregates was comparable between the two samples (Figure 1B-E). We then visualized clearance of aggresomes in the presence of CB-5083 (1 μM of CB-5083 was the highest tolerated dose in release experiments due to significant cell death at higher concentrations of the drug) and found an increase in the persistence of ubiquitin conjugates (Figure 1B, compare lane 5 and 6). This was confirmed in imaging studies where we find an increase in aggregates and to a lesser extent aggresomes, relative to untreated cells (Figure 1C, F and G). We conclude that long-term depletion of p97 (~72 hours in shRNA-based studies) results in the accumulation of various p97 substrates that are recruited to aggresomes increasing the size and percentage of aggresomes. However acute treatment (8 hours in CB-5083 studies) leads to direct inhibition of processes that recruit ubiquitylated substrates to aggresomes. Our studies suggest that the duration of p97 inhibition or depletion can have distinct outcomes that should be taken into account when designing studies to interfere with p97 function.

**Figure 1.**
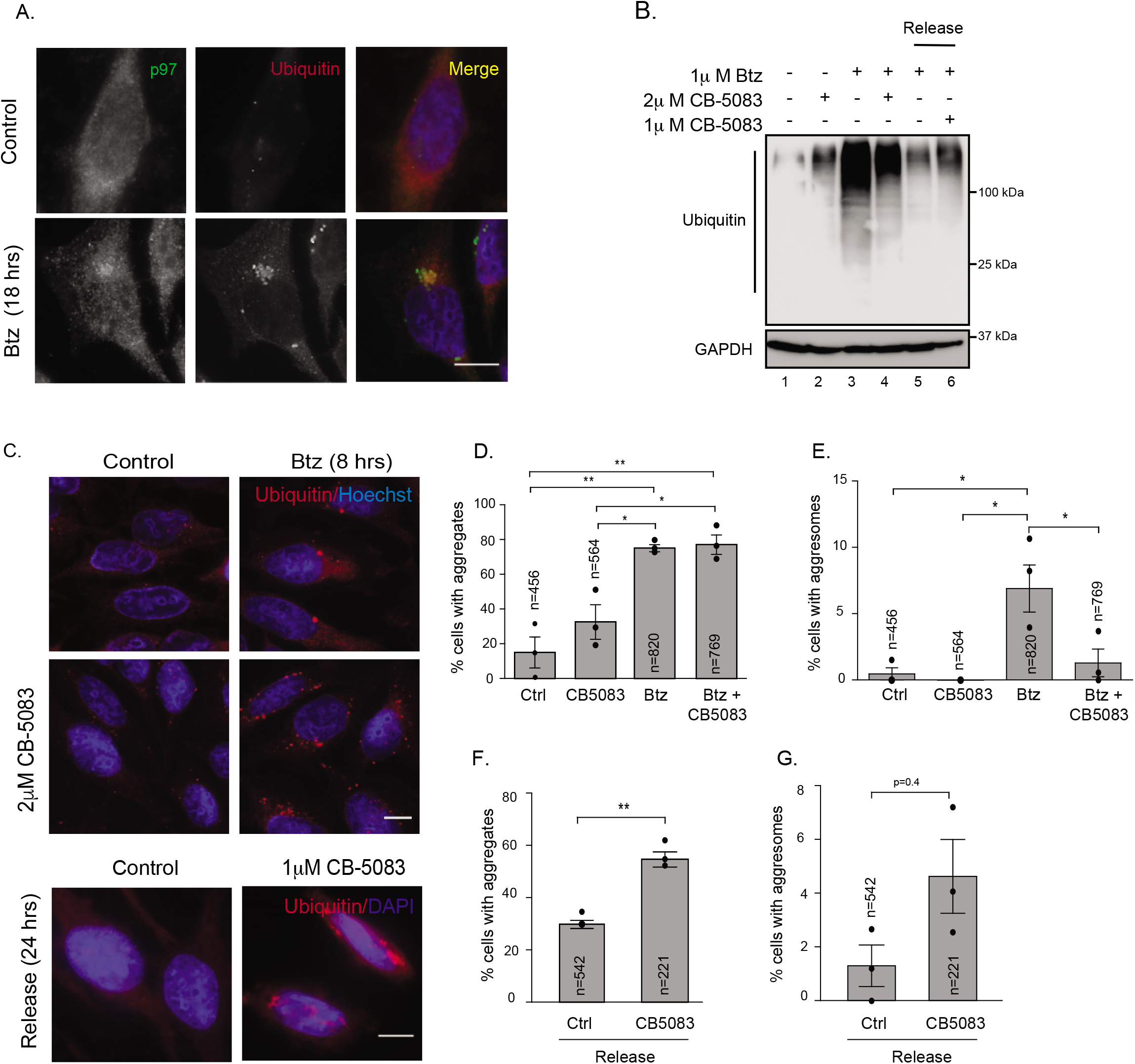
p97 is required for aggresome formation and clearance. (A) Hela Flp-in T-Rex cells were treated with 1 μM Btz for 18 hrs. Cells were stained for p97, ubiquitin (FK2), and nuclei (Hoechst dye). (B) Hela Flp-in T-Rex cells were treated with 1 μM Btz, 2 μM CB-5083, or both for 8 hrs. Cells were released into drug-free media or media containing 1 μM CB-5083 for 24 hours. Cell lysates were probed for ubiquitin. (C) Cells were treated as in (B) and imaged for ubiquitin-positive aggregates and aggresomes. (D) Total cellular aggregates (encompassing cytosolic and perinuclear aggregates) were quantified using AggreCount for images in (C). (E) Perinuclear aggresomes (minimum size cutoff 2 μm^2^) were quantified using AggreCount for the images in (C). (F) Total cellular aggregates (encompassing cytosolic and perinuclear aggregates) in the release samples were quantified using AggreCount for images in (C). (G) Perinuclear aggresomes in release samples were quantified using AggreCount for the images in (C). The indicated number of cells was analyzed from the three independent biological replicates indicated by the black dot. Graphs show the mean +/- s.e.m. *: p<=0.1, **: p<=0.05, ***: p<=0.001 as determined by One-way ANOVA with Bonferroni correction. Scale bar: 10μm.

### The p97 adaptors UBXN1 and NPL4 are localized to aggresomes

Given the significant role of p97 in aggresome formation and clearance, we next investigated the role of p97 adaptors in this process. We focused on the UBXD family, as these are the largest family of dedicated p97 adaptors, and many contain ubiquitin-associated domains (UBA) for associating with ubiquitin chains on substrates. Notably, p97 mutations known to cause MSP, have been reported to alter UBXD adaptor binding^28,29^. We also tested the involvement of the UFD1-NPL4 dimer as they participate in most known p97-dependent processes. We first screened for UBXD adaptors that localized to the aggresome upon proteasome inhibition using antibodies to endogenous proteins. Of the nine adaptors tested (seven UBXD proteins, UFD1 and NPL4), we found that UBXN1 was the only UBXD adaptor that robustly localized to the aggresome (Mander’s co-efficient of localization: 0.9 +/- 0.087, Figure 2A, Supplementary Figure 1A, Supplementary Figure 2A). Consistent with previous reports,^34^ we also observed recruitment of UFD1 and NPL4 to aggresomes (Mander’s co-efficient of localization: UFD1: 0.842 +/- 0.065, NPL4: 0.870+/- 0.073, Figure 2B). In contrast, the UBXD adaptor p47 was not present at aggresomes (Figure 2B). In time-course studies, we find that UBXN1 is recruited as early as 6 hours post-Btz treatment to aggresome-like structures as soon as ubiquitin staining is visible (Supplementary Figure 1A). We verified that UBXN1 structures were aggresomes by staining with canonical aggresome markers such as Proteostat, HDAC6, and the 20S proteasome (Figure 2C). In agreement with the dependence of aggresome formation on the microtubule network, nocodazole treatment caused the dispersal of aggresomes and numerous, UBXN1 and ubiquitin-positive foci were observed instead (Figure 2D). UBXN1 localization to aggresomes was not limited to HeLa cells but was observed in multiple cell types including the neuroblastoma cell line SH-SY5Y and the multiple myeloma cell line MM1.S (Supplementary Figure 2B and C). We next asked if the localization of UBXN1 to ubiquitin-positive aggregates was limited to those formed as a result of proteasome inhibition or if UBXN1 recognized a wider array of protein aggregates induced by distinct mechanisms. We found UBXN1 does not co-localize with stress granules formed as a result of sodium arsenite treatment (Figure 2E). Unexpectedly, we observe the recruitment of another p97 adaptor, p47 to stress granules; the significance of this is presently under investigation (Figure 2E). UBXN1 is not recruited to P-bodies that are sites of mRNA decay (Figure 2F). However, UBXN1 colocalization was observed with aggresome-like induced structures (ALIS) that contain defective ribosomal products that arise from treatment with translational poisons such as puromycin (Supplementary Figure 2D)^35^. Consistent with previous reports, we find that aggresome formation and poly-ubiquitin accumulation can be reversed by inhibiting protein translation with cycloheximide (Supplementary Figure 2E and F)^36^. Taken together, we have identified UBXN1 as an aggresome-localized protein that likely recognizes specific types of misfolded, ubiquitylated proteins.

**Figure 2.**
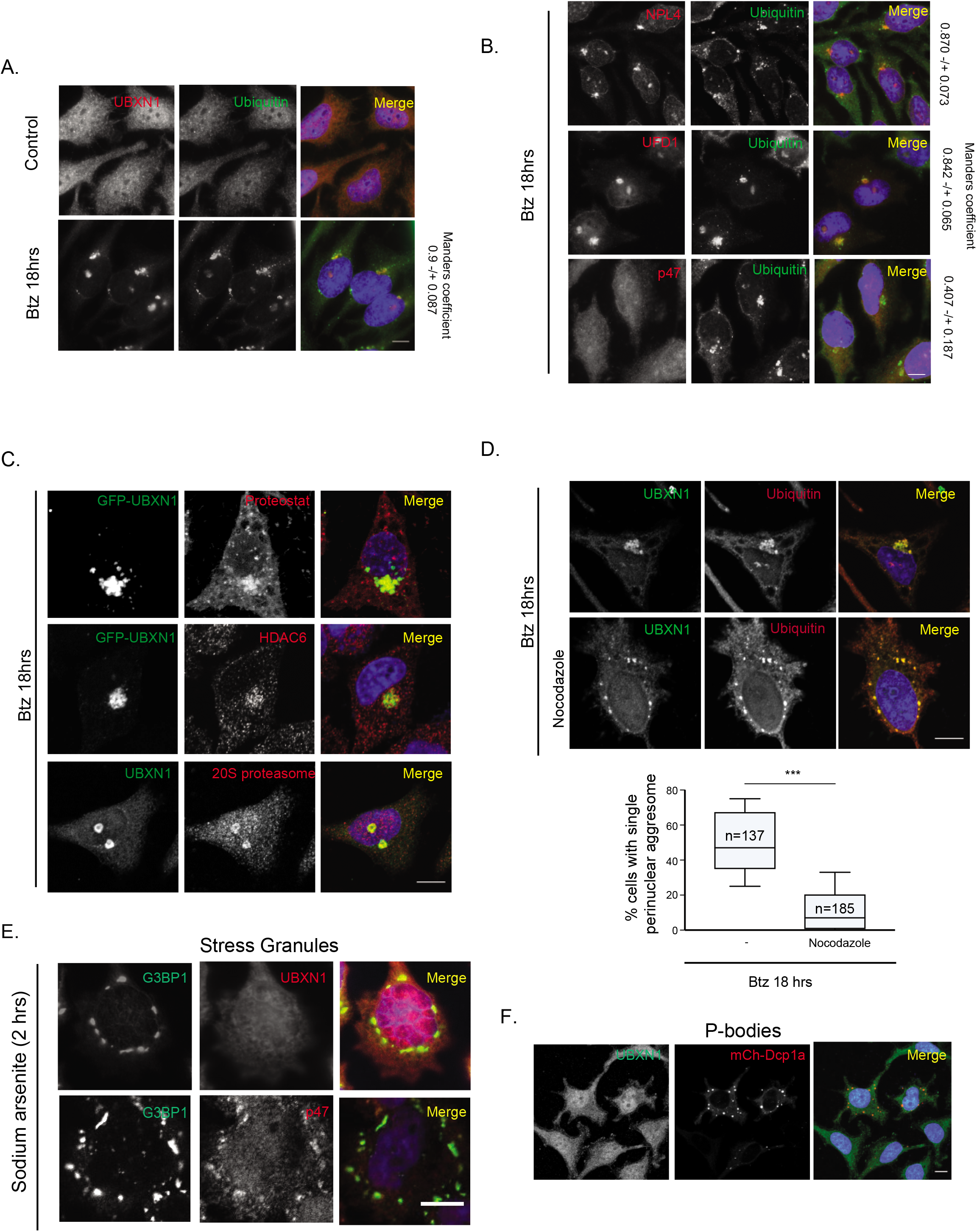
The p97 adaptors UBXN1 and NPL4 are localized to aggresomes. (A) Hela Flp-in T-Rex cells were treated with 1 μM Btz for 18 hrs and cells were stained for UBXN1 and ubiquitin. Co-localization was determined by the Mander’s overlap coefficient for 25 cells in 3 replicate experiments. (B) NPL4, UFD1 and p47 localization to aggresomes labeled with ubiquitin. Co-localization was determined by the Mander’s overlap coefficient for 25 cells in 3 replicate experiments. (C) GFP UBXN1 co-localizes with aggresome markers: Proteostat, HDAC6, and 20S proteasomes in Btz treated cells. (D) Microtubules are required for GFP-UBXN1 localization to aggresomes. Nocodazole co-incubation in Btz treated cells prevents aggresome formation. Lower panel: The number of aggresomes was quantified. (E) Hela Flp-in T-Rex cells were treated with 0.1 mM sodium arsenite for 2 h. Cells were stained for stress granule marker G3BP1 and p97 adaptors (UBXN1 or p47, used here as a positive control. (F) Stable mCherry-Dcp1a cells (labeling P bodies) were stained with UBXN1. The indicated number of cells was analyzed from the three independent biological replicates. Graphs show the mean +/- s.e.m. ***: p<=0.001 as determined by unpaired Students t-test. Scale bar: 10μm.

### UBXN1 is required for the formation of aggresomes

To determine the role of UBXN1 in aggresome formation, we used UBXN1 knock-out (KO) HeLa Flp-in TRex cell lines created using CRISPR-Cas9 gene-editing that we published previously (Figure 3A)^35^. We assessed aggresome formation in response to Btz treatment in two separate UBXN1 KO clonal lines. We find that loss of UBXN1 results in the loss of a single perinuclear aggresome and an increase in aggregates throughout the cytosol (Figure 3B and E, Supplementary Figure 3A). We additionally verified the UBXN1 KO phenotype by transient depletion of UBXN1 with two separate siRNAs and similarly observed the loss of aggresomes (Supplementary Figure 3B). Notably, there was no change in total ubiquitin conjugates between wildtype and UBXN1 KO cells (Figure 3C). To rule out off-target effects, we introduced doxycycline-inducible GFP-UBXN1 into the single FRT site in the UBXN1 KO cell line. Induction of GFP-UBXN1 reinstated aggresome formation as well as GFP-UBXN1 localization to these structures in response to Btz treatment (Figure 3A, D, and E). We examined the possibility of rapid onset and dissolution of aggresomes in KO cells that were missed in our endpoint assay and performed time-course studies in the UBXN1 KO cells treated with Btz. However, we were unable to observe aggresomes at all time points tested, ruling out temporal differences in aggresome formation in UBXN1 KO cells (Figure 3F).

**Figure 3.**
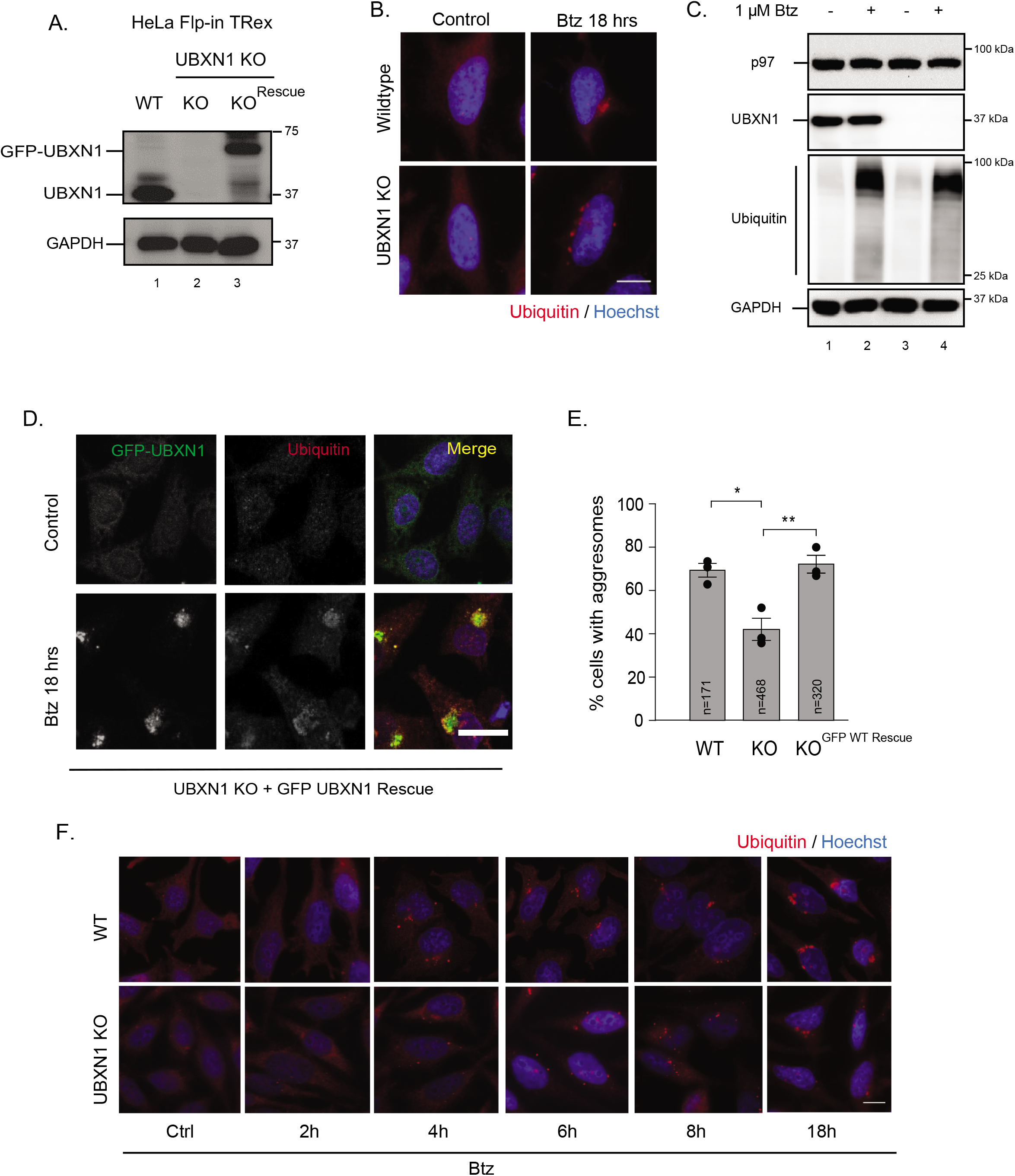
UBXN1 is required for the formation of aggresomes. (A) Immunoblot showing loss of UBXN1 in CRISPR-Cas9 generated knockout cells and re-expression of wildtype GFP-UBXN1 by doxycycline induction. (B) Wildtype (WT) and UBXN1 knock-out (KO) HeLa Flp-in TRex cell lines were treated with 1 μM Btz for 18 hrs. Cells were stained for ubiquitin. (C) Cell lysates from wildtype and KO celss treated with 1 μM Btz for 18 hrs were probed for ubiquitin. (D) GFP-UBXN1 expression in UBXN1 KO cells was induced by doxycycline. Cells were treated with Btz and stained for ubiquitin. Expression of GFP-UBXN1 reinstated aggresome formation in UBXN1 KO cells. (E) Quantification of data in panels (B and D). (F) WT and UBXN1 KO lines were treated with Btz for the indicated times and stained for ubiquitin. The indicated number of cells was analyzed from the three independent biological replicates indicated by the black dot. Graphs show the mean +/- s.e.m. *: p<=0.1, **: p<=0.05, as determined by One-way ANOVA with Bonferroni correction. Scale bar: 10μm.

While we did not observe other UBXD adaptors recruited to the aggresome, we nevertheless asked if other adaptors were required for the formation of aggresomes. We created a p47 KO cell line and a NPL4 knockdown cell line using CRISPR-Cas9 gene-editing in HeLa Flp-in TRex cells (Figure 4A). We used siRNA-mediated depletion of p47 and NPL4 to confirm our findings and to additionally examine the roles of UFD1 and FAF1 (Supplementary Figure 4A and B). In addition to UBXN1, both NPL4 and UFD1 (but not p47 or FAF1) were required for aggresome formation (Figure 4A-C and Supplementary Figure 4A and B). These findings are in agreement with prior studies that identified a role for the UFD1-NPL4 dimer in aggresome formation and are not all together surprising given the importance of UFD1-NPL4 in p97 activity. To determine whether UBXN1 and UFD1-NPL4 functioned together on a single p97 hexamer or if they were part of exclusive p97 complexes we performed two-step affinity purifications in cells transfected with FLAG-NPL4 and Myc-UBXN1. In the first step, FLAG-NPL4 associated complexes were affinity purified via the FLAG epitope, and bound proteins were eluted with FLAG peptide. Endogenous p97 in the eluate was purified with a p97 antibody and the presence of Myc-UBXN1 was determined by immunoblotting. As seen in Figure 4D, UBXN1 was bound to the same p97 hexamer as NLP4 and the interaction was modestly stimulated by Btz treatment (lanes 7, 8, 11, and 12). We next attempted the co-depletion of UBXN1 and UFD1-NPL4 to observe the effect on aggresome formation. However, the co-depletion of UBXN1 and UFD1 or NPL4 in (by dual siRNA transfection) combination with Btz resulted in cell death precluding further analysis. Collectively, our studies have uncovered a novel role for UBXN1 in mediating aggresome formation.

**Figure 4.**
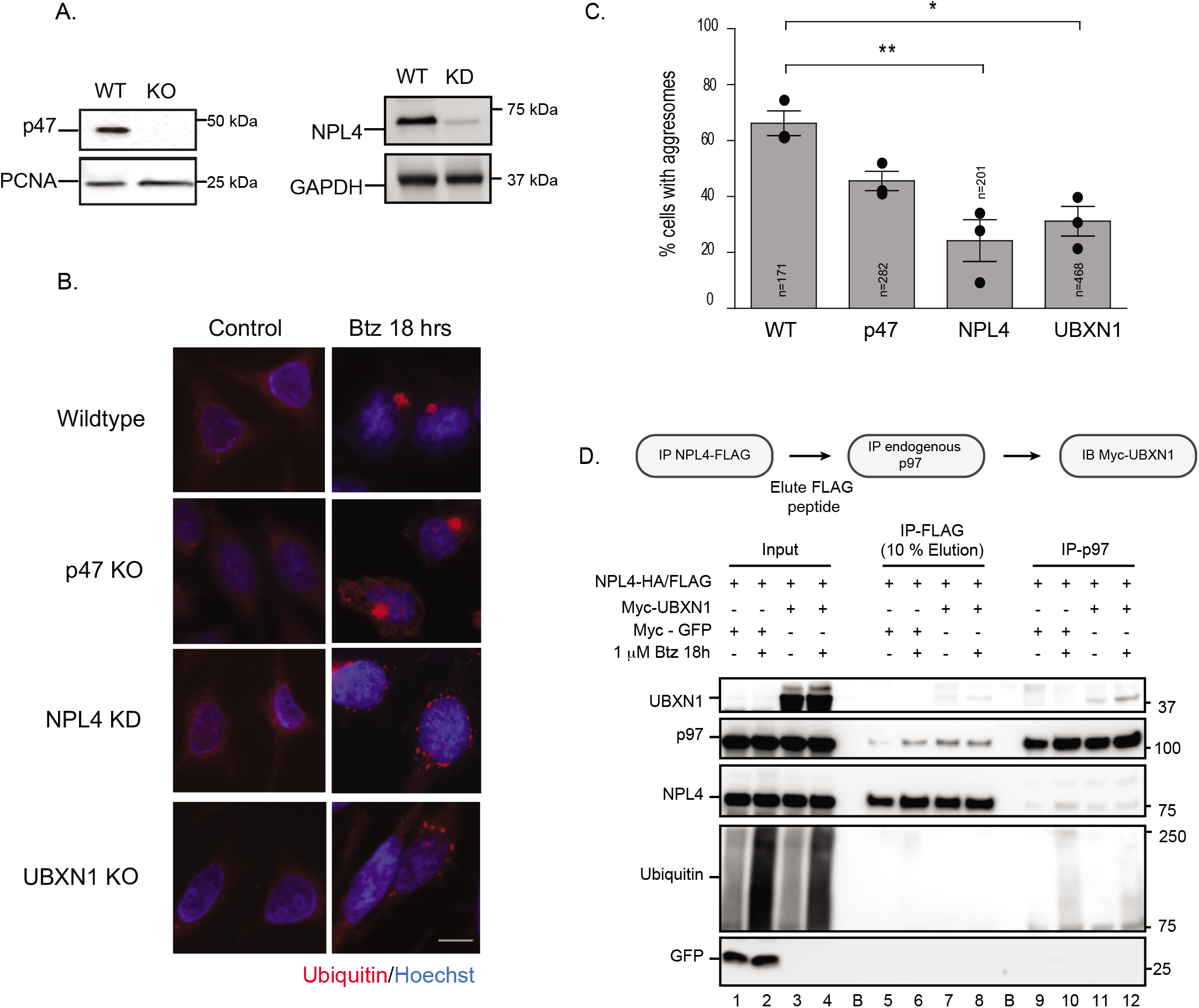
UBXN1 and NPL4 are required for aggresome formation and bind the same p97 molecule. (A) Immunoblot validation of CRISPR-Cas9 gene-edited knockout (p47) or knockdown (NPL4) cell lines. (B) Aggresome formation in CRISPR-Cas9 gene-edited knockout cell lines for the indicated UBXD adaptors. (C) Quantification of data in (B). (D) HEK293T cells were transfected with the indicated FLAG or Myc-tagged constructs. NPL4 was affinity purified using FLAG magnetic beads and associated proteins were eluted with FLAG peptide. Endogenous p97 in the eluate was immunopurified and probed for UBXN1, NPL4, and ubiquitin. UBXN1 associates with the same p97 hexamer associated with NPL4. The indicated number of cells was analyzed from the three independent biological replicates indicated by the black dot. Graphs show the mean +/- s.e.m. *: p<=0.1, **: p<=0.05 as determined by One-way ANOVA with Bonferroni correction. Scale bar: 10μm.

### p97 association is required for the role of UBXN1 in aggresome formation

Our results thus far suggest that p97 and its adaptors UBXN1 and UFD1-NPL4 are required for aggresome formation. Since the UFD1-NPL4 dimer functions in many p97-dependent processes, we focus on the role of UBXN1 in aggresome formation for the remainder of the study.

UBXN1 contains an N-terminal UBA domain that associates with ubiquitin and C-terminal ubiquitin-X (UBX) domain that interacts with p97, separated by a coiled-coil region (Figure 5A). We asked whether the association with ubiquitin and/ or p97 was required for aggresome formation. We mutated conserved residues within the UBA domain (Met^13^ and Phe^15^ to Ala, UBA^mut^) that mediate association with the hydrophobic isoleucine patch on ubiquitin or the UBX domain (Phe^265^Pro^266^Arg^267^ truncated to Ala-Gly, UBX^mut^) that associates with the N-terminus of p97. GFP-tagged mutants were introduced into the FRT site within the UBXN1 KO cell line and equal levels of induction of GFP-tagged proteins was determined by immunoblotting (Figure 5A and B). We verified that the mutants were unable to bind ubiquitin or p97 using co-immunoprecipitation assays in HEK293T cells expressing Myc-tagged UBXN1 constructs (wildtype, UBA^mut^ or UBX^mut^) and probing for endogenous ubiquitin or p97. As predicted, the UBXN1 UBA^mut^ failed to bind ubiquitin conjugates and UBXN1 UBX^mut^ was deficient in p97 binding (Supplementary Figure 5A and B). We next induced expression of GFP-UBXN1 wildtype, UBA^mut^, or UBX^mut^ in UBXN1 KO cells and assessed aggresome formation (based on perinuclear localization and size). Re-expression of wildtype UBXN1 rescued aggresome formation as before. However, UBX^mut^ expressing cells did not form aggresomes as well as the wildtype GFP-UBXN1; in these cells, the ubiquitin-positive aggregates were dispersed and fewer aggresomes were formed (Figure 5C-E). The UBXN1 UBA^mut^ also had modest defects in aggresome formation but not to the extent observed with the UBX^mut^ (Figure 5C-E, Supplementary Figure 5C). Notably, when we measured the fraction of cells that contained both cytosolic and perinuclear aggregates, it was comparable between wildtype and the two mutants, indicating that the aggregate burden was similar in all cases, ruling out increased clearance in the mutant expressing cells (Figure 5D). These results suggest that association of UBXN1 with p97 (and to a lesser extent ubiquitin) is an important requirement in aggresome formation. To determine whether deregulated aggresome dynamics resulting from p97, UBXN1, or NPL4 loss of function impacted cell survival in the face of proteasome impairment, we measured cell viability using a luminescence-based assay. We find that depletion of p97, UBXN1, and NPL4 sensitized cells to Btz treatment compared to wildtype cells (Figure 5F). Importantly, re-expression of wildtype UBXN1 but not UBA^mut^ or UBX^mut^ rescued survival to levels observed in wildtype cells treated with Btz (Figure 5F). Taken together, our results suggest that the distinct steps of aggresome formation and clearance are mediated by UBXN1, UFD-NPL4, and p97 respectively and that both these events are independently important for alleviating proteotoxic stress.

**Figure 5.**
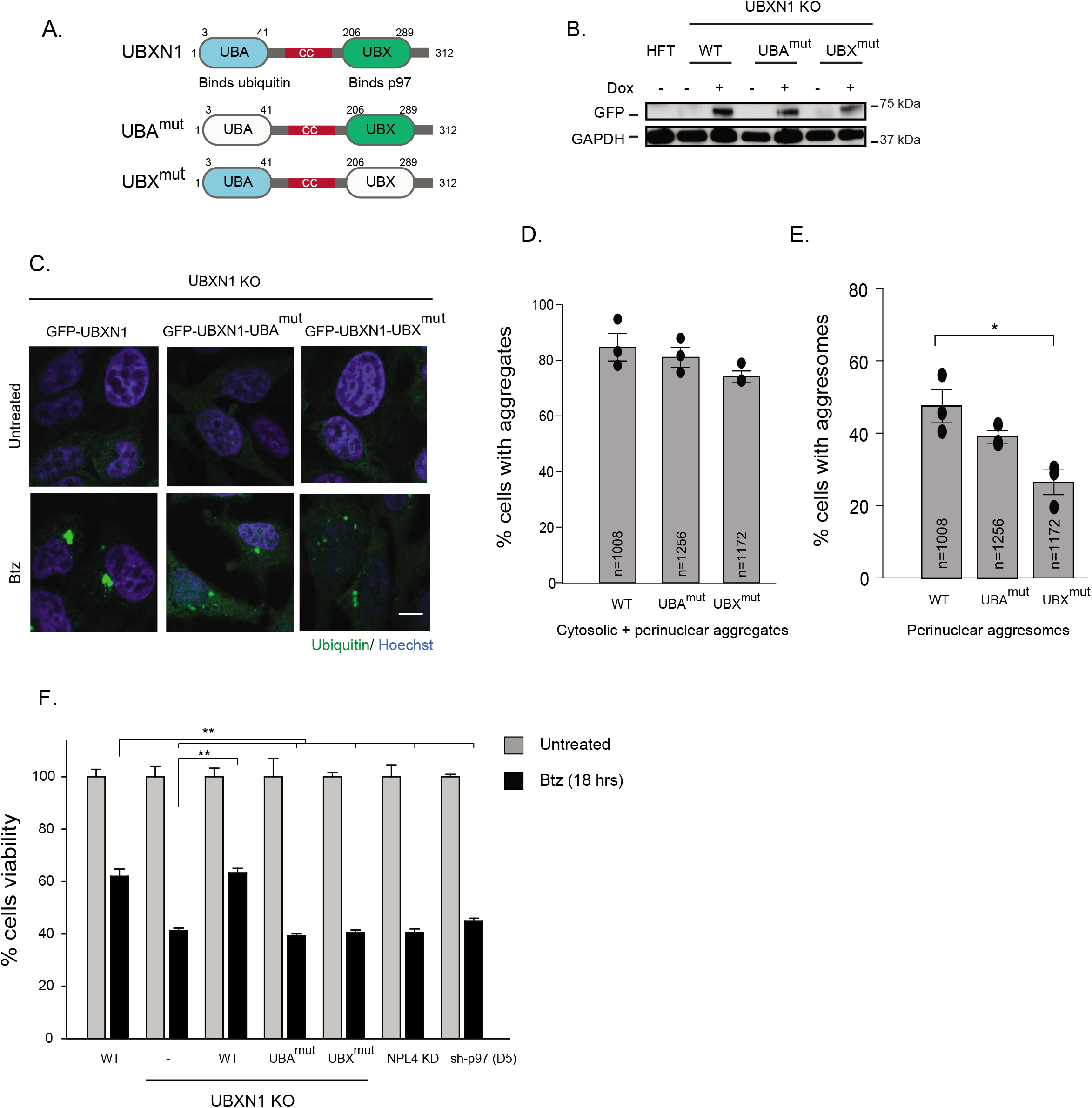
Analysis of domains within UBXN1 that are necessary for aggresome formation. (A) Domain organization of UBXN1 showing N-terminal ubiquitin associated domain (UBA), coiled coil domain (cc) and C-terminal ubiquitin X domain (UBX). (B) Expression of GFP-UBXN1, GFP-UBXN1-UBA^mut^ and GFP-UBXN1-UBX^mut^ in the UBXN1 KO cell line by the addition of doxycycline for 72 hrs. (C) UBXN1 KO cells expressing GFP-UBXN1 wildtype, GFP-UBXN1-UBA^mut^ or GFP-UBXN1-UBX^mut^ were treated with Btz for 18 hours and stained for ubiquitin. Re-expression of wildtype UBXN1 but not the UBX^mut^ rescued aggresome formation. The UBA^mut^ has smaller aggresomes but did not reach significance (D) Total cellular aggregates (encompassing cytosolic and perinuclear aggregates) were quantified for images in (C). (E) Perinuclear aggresomes were quantified for the images in (C). (F) Wildtype, UBXN1 KO (and indicated rescue lines), NPL4, and p97 depletion cell lines were plated in triplicate into 96-well plates and treated with 1 μM bortezomib for 18 hrs. Cell viability was measured and normalized to the value of untreated controls for each cell line. The indicated number of cells was analyzed from the three independent biological replicates indicated by the black dot. Graphs show the mean +/- s.e.m. *: p<=0.1, **: p<=0.05 as determined by One-way ANOVA with Bonferroni correction. Scale bar: 10μm.

Our studies indicate that UBXN1 acts early in the recognition of ubiquitylated substrates to mediate aggresome formation in collaboration with p97. Given that p97-depleted or inhibited cells were also unable to clear aggregates upon Btz removal, we asked if the same was true for the aggregates observed in cells lacking UBXN1. Surprisingly, we find that cells lacking UBXN1 or NPL4 were capable of clearing aggregates upon Btz removal (Supplementary Figure 5D). One possibility is that p97 utilizes distinct partners for the formation and clearance of aggresomes. p97 has been reported to associate with HDAC6 in prior studies and HDAC6 overexpression enhances clearance of aggregates in cells expressing p97 disease mutants^18^. In agreement with previous studies^17^, we find that p97 and HDAC6 associate with each other when transiently overexpressed in cells (Supplementary Figure 5E). The depletion of HDAC6 resulted in the loss of aggresomes as expected (Supplementary Figure 5F). Notably, HDAC6-depleted cells were less able to clear aggregates upon Btz removal (Supplementary Figure 5F). Taken together, our data suggest that p97-UBXN1 is required for the formation of aggregates, but that aggresome clearance is mediated via other p97 adaptors or interactors, possibly HDAC6.

### Loss of p97 and UBXN1 leads to an increase in Huntingtin polyQ aggregates

We were interested in determining whether UBXN1 has roles in the recognition of disease-relevant aggregates. Expansion of a CAG tract in the first exon of the huntingtin (htt) gene beyond a threshold of approximately 35-40 repeats causes Huntington’s disease (HD) and results in a mutant HTT protein containing an expanded polyglutamine (polyQ) segment^37^. Expression of exon 1 htt fragments longer than 40 residues leads to the formation of insoluble amyloid-like aggregates of HTT known as inclusion bodies^38–40^ (that are most reminiscent of aggresomes) and contributes to neurodegeneration. We used a previously published, doxycycline-inducible HTT polyQ91-mCherry U2OS cell line to investigate the role of UBXN1 in inclusion body formation^41^. Treatment with doxycycline leads to the appearance of ubiquitin and mCherry-Q91 positive aggregates in about 1-2% of cells consistent with previous studies^41,42^. A recent study suggested that the ubiquitylation of HTT inclusion bodies is not a pre-requisite for their formation. We used the ubiquitin E1 inhibitor TAK-273 at a dose that depleted endogenous ubiquitin conjugates in both untreated and Btz treated cells (Supplementary Figure 6A) and indeed found that Q91 inclusion bodies still formed but were not ubiquitylated (Supplementary Figure 6B). However, in cells treated with TAK-273, there were significantly more Q91 inclusion bodies compared to untreated cells, to a similar extent as Btz treatment (Supplementary Figure 6C). Unlike the role of p97 in aggresome formation (Figure 1), we find that inhibition of p97 using CB-5083 resulted in an increase in Q91 inclusion bodies (Supplementary Figure 6C). Thus, while the formation of HTT inclusion bodies does not require ubiquitylation, it is dependent on the ubiquitin-proteasome system and p97 for clearance.

We observed recruitment of UBXN1 to Q91 aggregates where it formed a ring-like structure around the aggregate, similar to ubiquitin (Figure 6A). We tested the impact of UBXN1 depletion on Q91 aggregate formation using siRNA transfection followed by microscopy. Surprisingly, loss of UBXN1 resulted in an increase in the number of polyQ aggregates in cells in a manner comparable to proteasome or p97 inhibition (Figure 6B and Supplementary Figure 6C and D). This was unexpected given the role of UBXN1 in aggresome formation. The high fluorescent intensity of Q91 aggregates necessitated lower capture settings during image acquisition that prevented us from quantifying smaller, less intense aggregates. To address this shortcoming, we used Pulse Shape Analysis (PulSA) by flow cytometry that has been used to analyze particles of different sizes including polyQ aggregates^41,42^. PulSA uses standard pulse width and height in the fluorescent channel. The formation of HTT containing aggregates leads to a decrease in pulse width and an increase in pulse height, allowing for the visualization of smaller aggregates and larger inclusion bodies. As we had observed from imaging studies, PulSA analysis of p97 or UBXN1 depleted cells indicated an approximately three-fold increase in polyQ inclusion bodies compared to control samples (Figure 6C). PulSA also enabled us to measure smaller aggregates that were missed in the imaging studies. We found an increase in smaller Q91-mCherry aggregates in cells depleted of UBXN1 or p97 suggesting that loss of this complex promotes the transition of Q91 oligomers into aggregates (Supplementary Figure 6E).

**Figure 6.**
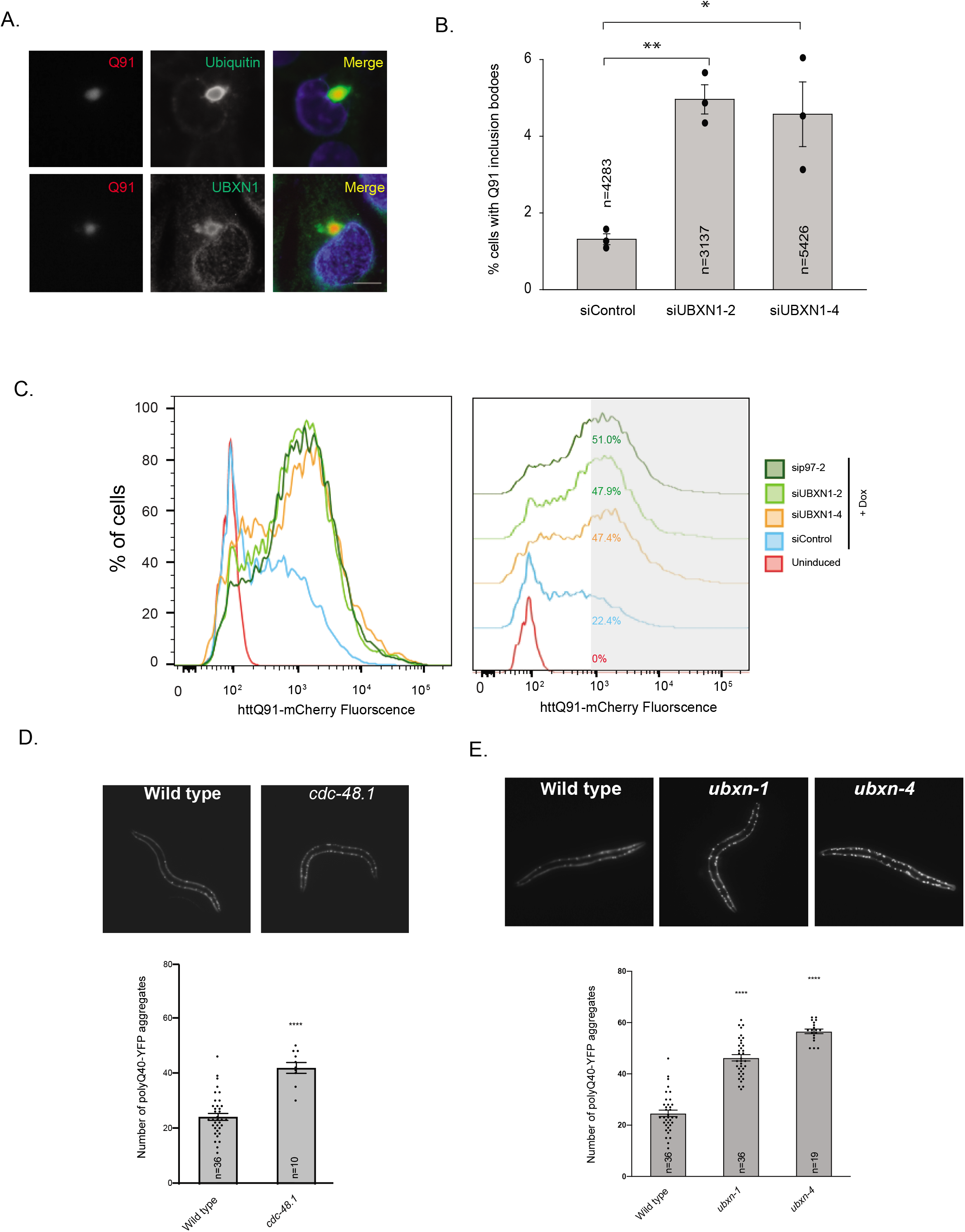
Role of p97 and UBXN1 in Huntingtin polyQ aggregation. (A) U2OS cells were treated with doxycycline to induce the expression of HTT Q91-mCherry. Cells were fixed and stained for ubiquitin and UBXN1 to demonstrate co-localization with HTT Q91-mCherry. (B) UBXN1 was transiently depleted with siRNAs in U2OS HTT Q91-mCherry and imaged for inclusion body formation. The number of HTT Q91-mCherry inclusion bodies was quantified. (C) PuLSA analysis of HTT Q91-mCherry aggregates in UBXN1 and p97 depleted cells. (D and E) Representative fluorescent images of L4 larvae stage wild type, *cdc-48.1(tm544) (D), ubxn-1(tm2759)* and *ubxn-4(ok3343)* (E) loss-of-function mutants expressing polyQ40::YFP in body wall muscle. Images were taken of worms precisely age-matched at the L4.4 vulva developmental stage. Bottom panels in each show quantification of visible fluorescent aggregates in L4 larvae animals expressing polyQ40::YFP in either wildtype, *cdc-48.1, ubxn-1* and *ubxn-4* mutant animals. Quantification was only performed on worms at the L4.4 vulva development stage based on vulva morphology. The indicated number of cells was analyzed from the three independent biological replicates indicated by the black dot. Graphs show mean +/- s.e.m. *: p<=0.1, **: p<=0.05, ***: p<=0.001 as determined by One-way ANOVA with Tukey (B) or Dunnett’s Test (D and E). Scale bar: 10μm.

We next asked if p97-UBXN1 regulated polyQ aggregation in vivo. We used transgenic worms expressing YFP-polyQ40 under the control of the *unc-54* myosin heavy chain promoter to direct expression in the body wall muscle cells (mQ40)^43^. Previous studies have demonstrated that mQ40 marks a threshold at which the reporter begins to aggregate in worms^43^. There are two isoforms of p97 (*cdc*-48.1 and *cdc*-48.2) with redundant functions in worms. In addition, six UBXD adaptors (including *ubxn*-1) and *ufd*-1 and *npl*-4 are conserved. We crossed *cdc*-48.1, *ubxn*-1, and *ubxn*-4 (worm ortholog of mammalian UBXD2) mutant worms with the mQ40 strain. To accurately quantify aggregates in control and mutant strains, worms were age-matched at the L4.4 vulval development stage, and aggregates were quantified by microscopy. We found that control mQ40 worms typically had on average 20 aggregates, however *cdc*-48.1 and *ubxn*-1 mutant worms had a two-fold increase in mQ40 aggregates in agreement with our findings in mammalian cells (Figure 6D and E). This finding is in agreement with a previous study where the over-expression of either *cdc-48* isoform in mQ40 expressing worms partially suppresses Q40 aggregation^44^. Surprisingly, *ubxn-4* mutant worms also displayed an increase in mQ40 aggregates, indicating that additional adaptors may be utilized for HTT aggregate clearance in worms (see Discussion).

In summary, we have identified an evolutionarily conserved role for p97-UBXN1 in the recognition, formation, and clearance of ubiquitylated protein aggregates.

## Discussion

Cellular proteostasis networks are complex yet flexible to accommodate the need for thousands of proteins to perform various cellular functions while minimizing protein misfolding and aggregation. Defects in this large network result in multi-system proteinopathies, and several recent reports suggest that the age-related decline in the quantity, capacity, and efficiency of these systems may be an underlying cause of these disorders^7^. Whether aggregates observed in human diseases are causative of disease-relevant phenotypes or if they represent a cytoprotective structure meant to sequester and shield aggregates is controversial.

However, at least in the case of polyQ inclusions observed in Huntingtin’s disease, the soluble microaggregates are believed to be more toxic than the inclusion body^14,45,46^.

In this study, we have identified a novel role for the p97 adaptor UBXN1 in the recognition of specific types of cellular aggregates. UBXN1 is the only UBXD adaptor that robustly localizes to aggresomes and cells lacking UBXN1 have a diminished ability to form a single aggresome but instead display an increased number of dispersed microaggregates. In particular, the total ubiquitin burden (measured by immunoblotting total cellular ubiquitin conjugates as well as quantifying cytosolic and perinuclear aggregates by imaging) remains comparable in wildtype and UBXN1 KO cells. While p97 has been implicated in a large number of ubiquitin-dependent events, few of its numerous adaptors have been linked to these processes. Indeed, apart from the UFD1-NPL4 dimer that functions in most p97 processes (due to dual roles in enhancing ubiquitylated substrate binding to p97 as well as stimulating p97’s unfoldase activity)^20,47^ many of its adaptors including UBXN1 remain poorly studied. A single UFD1-NPL4 dimer is believed to associate with the p97 hexamer, and at least in the case of FAF1 and UBXD7, UFD1-NPL4 association with the p97 hexamer is a pre-requisite for binding^48^. Our studies suggest that a fraction of the cellular pool of UBXN1 and the UFD1-NPL4 dimer associate with the same p97 hexamer. The presence of dual adaptors on p97 may increase the capacity of a single hexamer in capturing multiple substrates or may enhance the avidity of substrate interaction by providing multiple ubiquitin-binding domains on the same molecule. Surprisingly, neither UBXN1 nor NPL4 are required for aggresome clearance following Btz removal and suggests that p97 may switch adaptors for clearance purposes. In support of this, we and others have found that p97 interacts with HDAC6, which is required for aggresome clearance and may collaborate with p97 in this process. Identifying sites of interaction between these molecules and testing interaction-deficient mutants in aggresome formation will provide conclusive evidence for the role of a p97-HDAC6 complex in aggresome clearance. What factors may dictate adaptor choice during these distinct phases remains to be determined.

Surprisingly, depleting p97 protein levels or inhibiting its catalytic activity have distinct outcomes in terms of aggresome formation. We find that long term depletion of p97 leads to an increase in cells with aggresomes that are larger compared to control cells. In contrast, acute p97 inhibition with CB-5083 leads to loss of aggresomes. Previous studies using ubiquitin remnant capture proteomics suggest that while many p97 and 26S proteasome targets are shared, p97 additionally processes ubiquitylated substrates that are not proteasomal targets^49^. Thus, the long-term accumulation of these substrates may contribute to the enhanced aggresome phenotype. In contrast, both depletion and inhibition of p97 lead to defects in the clearance of aggresomes.

We find that depletion of p97 and UBXN1 increases polyQ-containing aggregates in mammalian cells in vitro and in a *C.elegans* model of Huntington’s disease. Using PuLSA, we show that depletion of p97 and UBXN1 in mammalian cells leads to an increase in smaller aggregates reminiscent of the aggresome phenotype. However, we also observe an increase in large inclusion bodies, which is distinct from the role of p97-UBXN1 in aggresome formation. What factors could account for these different phenotypes? Inclusion bodies share several features with aggresomes; they are ubiquitylated, frequently (though not exclusively) form near the perinuclear area, are dependent on microtubules for their formation, and recruit similar protein quality control components^50^. However, there may be distinctions that influence what proteins are recruited to each structure and the discrete steps that culminate in their removal. Aggresomes are composed of a diverse array of ubiquitylated proteasomal substrates, whereas inclusion bodies are seeded by misfolded HTT that eventually trap other cellular components. UBXN1 may implement aggresome formation via association with a distinct cohort of proteins (both ubiquitylated substrates and regulators). However, the increase in smaller polyQ aggregates in UBXN1-depleted cells may enable direct seeding of larger inclusion bodies. We have previously shown that loss of UBXN1 impairs the clearance of ALIS^35^ whereas UBXN1 appears to have no role in aggresome clearance further underscoring distinctions in how different aggregates are handled by the same protein. In worms, *ubxn*-1 does not appear to be the sole p97 adaptor in the recognition of inclusion bodies, we find that *ubxn*-4 mutant worms also have increased polyQ aggregates. Mammalian UBXD2 (ortholog of *ubxn*-4) is a component of ERAD and the increased polyQ aggregation that we observe in *ubxn*-4 may be due to inhibition of ERAD and the accumulation of misfolded proteins that exacerbates polyQ aggregation. Further studies are warranted to explore the role of other p97 adaptors in polyQ aggregation.

How does UBXN1 recognize cargo destined for the aggresome? One possibility is the recognition of ubiquitin chains on substrates by the UBA domain in UBXN1. However, our studies with UBA and UBX mutants suggest that while ubiquitin binding is important, it is not as critical as p97 interaction. This suggests that additional features of aggresome-destined substrates may be directly or indirectly sensed by UBXN1. Recently, components of the linear ubiquitin chain assembly complex (LUBAC) E3 ligase were shown to be recruited to and modify HTT polyQ inclusion bodies with linear ubiquitin chains^51^. LUBAC recruitment was dependent on p97, and both p97 recruitment and LUBAC-mediated linear ubiquitylation of HTT polyQ aggregates were required for efficient clearance. What other protein quality control components associate with UBXN1 and the types of ubiquitin linkages are recognized by UBXN1 and will be a topic of future studies.

Aggresomes were identified more than twenty years ago^8^, yet the full complement of proteins that are recruited to aggresomes remains incomplete. It is unclear what distinguishes different types of protein aggregates from one another and how specific components of protein quality control pathways are appropriately engaged. For example, aggresomes have long been known to be encased in intermediate filament vimentin cages. Yet it was only recently that it was demonstrated that vimentin specified the recruitment of proteasomes to aggresomes and is an important determinant in how aggregates are triaged as adult neural stem cells exit quiescence^52^. Given the prevalence of protein aggregates in numerous neurodegenerative diseases, a complete understanding of the principles that govern aggresome formation and clearance will aid in our understanding of disease-relevant aggregates. Our study demonstrates that the process of aggresome formation and clearance are distinct events mediated by multiple proteins. Given the role of p97 mutations in human disorders related to aggregate clearance, it will be of interest to determine whether UBXN1 regulates their formation and if targeting this event may have therapeutic benefit.

## Materials and Methods

### Cell lines and culture

HEK-293T (ATCC), HeLa Flp-in TRex (a kind gift from Brian Raught, University of Toronto), SHSY-5Y (a kind gift from Wade Harper, Harvard Medical School), and U2OS HTTQ91-mCHERRY (a kind gift from Ron Kopito, Stanford University) cells were cultured in Dulbecco’s modified Eagle’s medium (DMEM) supplemented with 10% fetal bovine serum (FBS) and 100 U/ml penicillin and streptomycin. Multiple Myeloma MM1.S cells (gift from Filemon Dela Cruz, Memorial Sloan Kettering), were cultured in RPMI-1640 containing 10% FBS and 100U/ml penicillin and streptomycin. Cells are tested for mycoplasma contamination on a routine basis. Cells were maintained in a humidified, 5% CO_2_ atmosphere at 37°C.

### Antibodies, chemicals, and reagents

The rabbit p97 antibody used for immunofluorescence (10736-AP), UBXN1 (16135-1-AP), UFD1 (10615-1-AP), NPL4 (11638-1-AP), FAF1 (10271-1-AP), UBXD1 (14706-1-AP), UBXD2 (21052-1-AP), UBXD3 (26062-1-AP), UBXD5 (13109-1-AP), p47 (15620-1-AP) antibodies are from Proteintech. Rabbit p97 antibody for immunopurification and immunoblotting is from Bethyl (A300-589A), mouse FLAG-M2 antibody was from Sigma; mouse ubiquitin (FK2) antibody used for immunofluorescence was from EMD Millipore; ubiquitin antibody (clone P4D1) for immuno-blotting, GAPDH (sc-47724), and mouse Myc antibody was from Santa Cruz (sc-40). Rabbit HDAC6 antibody (clone D2E5) was from Cell Signaling Technologies (7558), 20S proteasome antibody (MCP257) was from EMD Millipore. Bortezomib and TAK273 were from Selleckchem, puromycin, doxycycline, cycloheximide, and nocodazole were from Sigma. Anti-FLAG magnetic beads were from Sigma and Myc magnetic beads were from Pierce.

### siRNA transfections and generation of CRISPR-Cas9 knockout cell lines

pTRIPZ sh-p97 hairpin vectors were purchased from Open Biosystems (clone D1: V2THS_111071 Clone D5: V2THS_201657). For the generation of shp97 stable cell lines, lentivirus was produced and packaged in HEK293T cells reported previously and subsequently used to infect HeLa Flp-in TRex cells. Stable cell lines were selected with 1 μg/ml puromycin^23^. The depletion of p97 was achieved by treatment with 4 μg/ml of doxycycline for the 24-72 hrs.

siRNAs targeting p97 (15975124), UFD1 (4392420), NPL4 (4392420), HDAC6 (4427038) were purchased from Ambion. siRNAs targeting UBXN1 (D-008652-01, D-008652-02), FAF1 (D-009106-03), p47 (D-017222-01) were purchased from or Dharmacon. For siRNA transfections, cells were trypsinized and reverse transfected with 30 nM siRNA by using Lipofectamine 3000 (Thermo Scientific) according to the manufacturer’s protocol in 6 well plates. These cells were then trypsinized and plated on 12 well plates with glass coverslips where appropriate. In general, cells were harvested at 60 hrs post-transfection.

The CRISPR-Cas9 gene-editing system was used to generate knockout cell lines in HeLa Flp-in T-REX cells. The guide sequences for UBXN1 (5’GCCGTCCCAGGATATGTCCAA3’), p47 (5’AGATCATCCACCAGCTCGTT3’), and NPL4 (5’GATCCGCTTCACTCCATCCG3’) were cloned into the pX459 vector carrying hSpCas9 and transiently transfected into HeLa cells by using Lipofectamine 3000 (Thermo Scientific) according to the manufacturer’s protocol. At 36 hrs post-transfection, the cells were pulsed briefly with 1 μg/ml puromycin for a further 24 hrs. The surviving cells were then serially diluted to achieve 1 cell / well in 96-well plates for clonal selection. The knockout of the gene of interest was verified by immunoblotting. HttQ91-mCherry expression was induced by treating cells with 1 μg/ml of doxycycline for 48 hours.

### Cloning and generation of inducible rescue lines

UBA and UBX point mutants in UBXN1 were created by overlap PCR and Gibson cloning into pDON223. Gateway recombination (Thermo Scientific) was used to recombine into either the pcDNA FRT/TO-GFP or the pDEST N-Myc vector. All clones were verified by Sanger sequencing. GFP-UBXN1 rescue cell lines were generated by transfecting pcDNA FRT/TO-GFP-UBXN1, UBA^mut^, or UBX^mut^ mutants along with pOG-44 (1:9 ratio) into UBXN1 KO cells by using Lipofectamine 3000. Stable integrant cells were treated with 200 μg/ml hygromycin for 7-10 days until visible colonies were observed. GFP-UBXN1 wildtype expression was induced with 1 μg/ml doxycycline whereas UBA^mut^ and UBX^mut^ were induced with 1.5 μg/ml doxycycline to achieve equal levels of expression for 72 hours.

### Cell lysis, transfections, and immunoprecipitation

Cells were lysed in mammalian cell lysis buffer (50 mM Tris-Cl [pH 6.8], 150 mM NaCl, 0.5 % Nonidet P-40, Halt protease inhibitors [Pierce], and 1 mM DTT). To recover total protein (soluble and insoluble), sodium dodecyl sulfate (SDS) was added to a final concentration of 1% (Figure 1B, Supplementary Figure 2F and 6A). Cells were incubated at 4°C for 10 min and then centrifuged at 14,000 rpm for 15 min. The supernatant was removed, and the protein concentration was estimated using the DCA assay (Bio-Rad).

Myc or FLAG magnetic beads in a 1:1 slurry in mammalian cell lysis buffer was added to samples and rotated at 4°C overnight. A magnetic stand (Thermo Fisher) was used to wash the beads three times with lysis buffer. Beads were resuspended in 25 μl SDS sample buffer, boiled briefly, and processed for SDS-PAGE and immunoblotting. FLAG elutions were performed two times with the 250μg/ml 3XFLAG peptide (ApexBio) for 30 minutes at room temperature. Endogenous p97 was further immunoprecipitated (Bethyl) at 4°C overnight. Beads were washed 3 times in lysis buffer and resuspended in Laemmli buffer and boiled briefly.

### Immunofluorescence and microscopy

Cells were grown on coverslips (no. 1.5) in a 12-well plate treated with 1 μM bortezomib (Btz), 2 μM TAK243, 2 μM CB5083 (or 1 μM CB5083 for release experiments), 100 μg/ml cycloheximide or 5 μg/ml nocodazole (for the last 4 hours of Btz treatment) for the indicated times. Cells were washed briefly in phosphate-buffered saline (PBS) and fixed with 4% paraformaldehyde/PBS at room temperature for 15 min or ice-cold methanol for 10 minutes at −20°C. Cells were washed in PBS then blocked in 2 % bovine serum albumin (BSA) 0.3 % TritonX-100 in PBS for 1 h. The coverslips were incubated with the indicated antibodies in a humidified chamber overnight at 4°C. Coverslips were washed and incubated with the appropriate Alexa Fluor-conjugated secondary antibodies (Molecular Probes) for 1 h in the dark at room temperature. Cells were washed with PBS, and nuclei were stained with Hoechst dye and mounted onto slides. All images were collected using a Nikon A1R scan head with a spectral detector and resonant scanners on a Ti-E motorized inverted microscope equipped with a 60XPlan Apo 1.4-numerical-aperture (NA) objective lens. The indicated fluorophores were excited with either a 405-nm, 488-nm, or 594-nm laser line. Images were analyzed by using FIJI (https://imagej.net/Fiji). Colocalization analysis was performed by using the Coloc2 and JACoP plug-ins in FIJI.

### PuLSA analysis of HTT Q91-mCherry aggregates

U2OS-HTTQ91mCherry cells were trypsinized and reverse transfected with 20 nM siRNA by using Lipofectamine 3000 (Thermo Scientific) according to the manufacturer’s protocol in 6 well plates. Expression of HttQ91-mCherry was induced with 1 μg /ml dox 8 hours after siRNA transfection, and cells were incubated for an additional 48 h before harvesting and analysis. For PulSA flow cytometric analysis, cells were analyzed on a flow cytometer (LSR II; BD) equipped with 535-nm lasers. Measurements for mCherry peak width, peak height, and total intensity was collected. Flow cytometry analysis software (Flow Jo; Tree Star) was used to analyze the fraction of cells with Inclusion bodies (IBs) by PulSA. Cells with IBs were identified and gated using a mCherry peak width versus peak height scatter plot. The percentage of cells was plotted against total mCherry fluorescence intensity. 10,000 cells were analyzed for each of the three independent experiments.

### Fluorescence microscopy and quantification of aggregates in C.elegans

The following strains were used in this study: N2 (Bristol), *rmIs133* [*unc-54p*::Q40::YFP] (polyQ40::YFP expressed under a muscle-specific promoter)^43^, *cdc-48.1(tm544), ubxn-1(tm2759)* and *ubxn-4(ok3343*). All strains were maintained at 20°C as described previously^53^.

Representative fluorescent images of polyQ40::YFP in L4 larval stage *C. elegans* were acquired using a Carl Zeiss Axiovert M1 microscope equipped with a 5X objective and a YFP filter (Zeiss). Images were collected with an Orca-ER charged coupled device (CCD) camera (Hamamatsu) and Metamorph (version 7.1) software. Exposure settings and gain were adjusted to fill the 12-bit dynamic range without saturation and were identical for all images. Larval-stage 4 (L4) animals were immobilized in a droplet of M9 containing 2.3mM levamisole (Tetramisole, Sigma) and placed on a 2% agarose pad containing 3.4 mM levamisole. To quantify fluorescent puncta, worms were precisely age-matched at the L4.4 vulva development stage based on vulva morphologyusing a 100X objective. Then the number of polyQ40::YFP aggregates were visually counted on the microscope using the 5X objective and recorded. All genotypes were blinded prior to the experiment. At least 3-5 animals were measured for each genotype across 3 independent experiments and statistical analyses were performed by ANOVA followed by Dunnett’s multiple comparisons test.

### Aggresome Quantification

Aggresomes were quantified using AggreCount^33^, an ImageJ macro we developed for the unbiased analysis of cellular aggregates. AggreCount can analyze hundreds of cells and images in a matter of minutes and provides quantification of number of aggregates, size of aggregates, and their cellular distribution (cytosolic, perinuclear, and nuclear) at single-cell resolution. We used a 2 μm^2^ minimum size cut-off for aggresomes formed after 8 hours of Btz treatment and a 4 μm^2^ minimum size cut-off for aggresomes formed after 18 hours of Btz treatment based on the mean size of the largest perinuclear aggregate from hundreds of analyzed cells (see quantification in Supplementary Figure 1B)

### Statistical Analysis

Several hundred cells (indicated in the figures) were counted for each of three biological replicates. Means and standard deviations or standard error of the mean for triplicate measurements were calculated, and statistical significance was calculated by using an unpaired Student t-test or One-Way ANOVA with the indicated post-hoc testing.

## Acknowledgments

We are grateful to Rakesh Ganji for creating the knock-out cell lines, members of the Raman lab, Shireen Sarraf, and Karl Munger for critical reading of the manuscript. S.M and M.R conceived the study, performed the experiments, and co-wrote the manuscript. J.K developed AggreCount and performed all image analysis, B.O and P.J performed the *C.elegans* studies. B.O was supported in part by 2R25GM066567 as a Post-Baccalaureate Research Program (PREP) trainee. This work is supported in part by the American Cancer Society Research Scholar grant (RSG-19-022-01-CSM) to M.R., National Institutes of Health grant (R01-GM127557) to M.R., (R21NS101534) to P.J., and Tufts Russo Family Award to M.R and P.J. We thank the *Caenorhabditis* Genetics Center (funded by the National Institutes of Health Office of Research Infrastructure Programs Grant P40ODD010440) and Shohei Mitani (National Bioresource Project) for *C. elegans* strains.

---

**Supplementary Figure 1.**
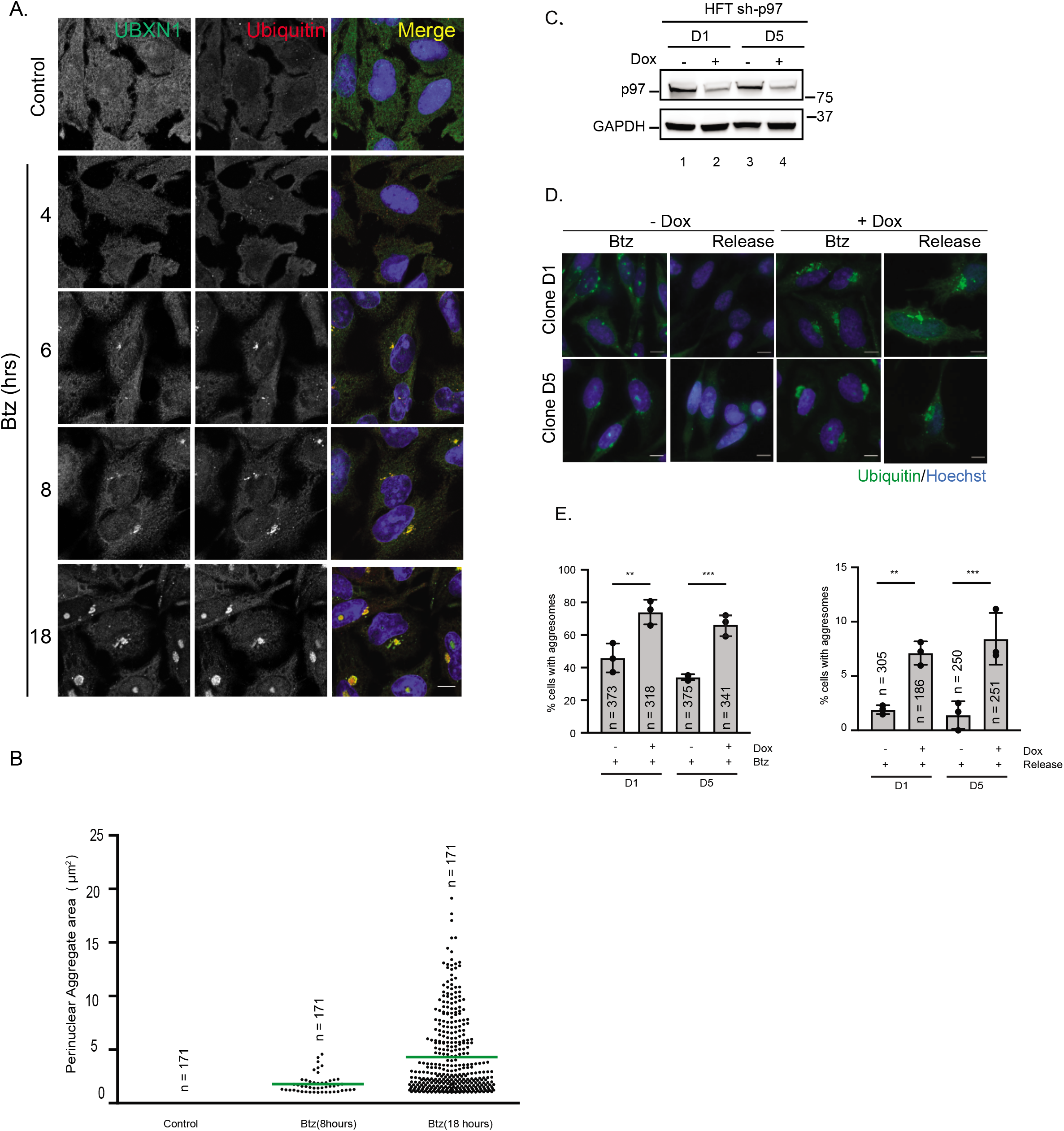
p97 is required for aggresome clearance. (A) Hela Flp-in T-Rex cells were treated with 1 μM Btz for 0, 4, 6, 8, or 18 hrs. Cells were stained with UBXN1 and ubiquitin and imaged (B) Hela Flp-in T-Rex cells were treated with 1 μM Btz for 0, 8, or 18 hrs. Cells were stained with ubiquitin and perinuclear aggregate size was quantified using AggreCount. The mean area of the largest perinuclear aggregate for each time point (2μm^2^ or 4μm^2^ respectively) was used for all subsequent analysis to quantify perinuclear aggresomes. (C) Levels of p97 depletion in doxycycline-inducible shRNA Hela Flp-in T-Rex cell lines. Cells were treated with doxycycline for 72 hours. (D) sh-p97 cell lines were treated with 1 μM Btz for 18hrs and released into drug-free media for a further 24 hours in the presence or absence of p97 depletion. Cells were stained with ubiquitin and Hoechst. The number of cells containing ubiquitin-positive aggresomes was quantified using AggreCount. (E) Quantification of data in (D). The indicated number of cells was analyzed from the three independent biological replicates indicated by the black dot. Graphs show mean +/- standard deviation. *: p<=0.1, **: p<=0.05, ***: p<=0.001 as determined by One-way ANOVA with Bonferroni correction. Scale bar: 10μm.

**Supplementary Figure 2.**
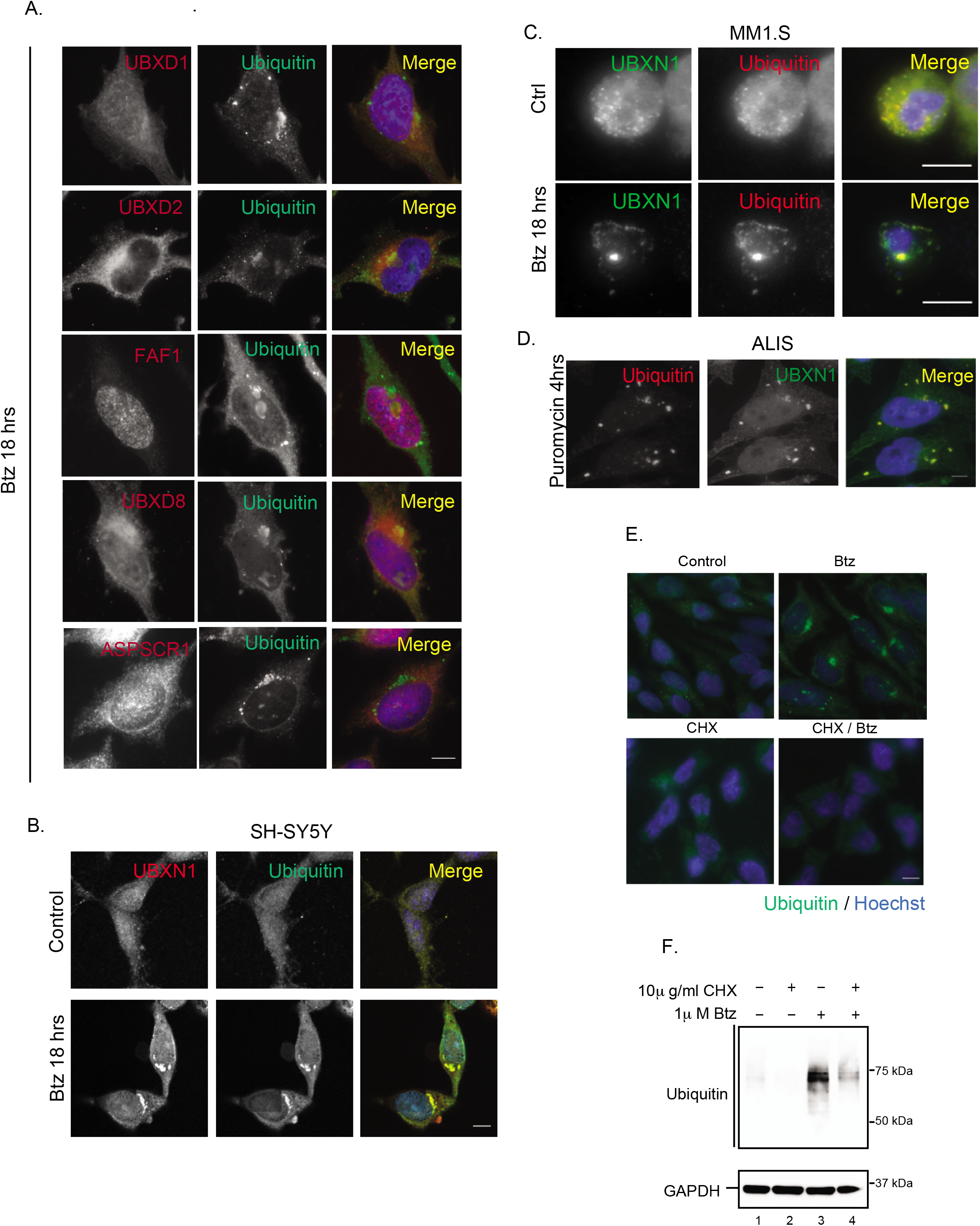
The p97 adaptors UBXN1 and NPL4 are localized to aggresomes. (A) HeLa Flp-in TRex cells were treated with 1 μM Btz for 18hrs. Cells were stained for p97 adaptors (UBXD1, UBXD2, FAF1, UBXD8, and ASPSCR1) and ubiquitin. Note that UBXD2 and UBXD8 are known to be ER-tethered adaptors and localize to both ER tubules at the cell periphery and ER sheets near the nuclear envelope. They do not co-localize with aggresomes and can be seen to occupy a greater cellular area (corresponding to ER sheets) than the aggresome. (B) Neuroblastoma cells SH-SY5Y were treated with 1 μM Btz for 18 hrs. Cells were stained for UBXN1 and ubiquitin. (C) Multiple Myeloma cell line MM1.S was treated with 1 μM Btz for 18 hrs. Cells were stained for UBXN1 and ubiquitin. (D) Hela Flp-in T-Rex cells were treated with 5 μg/ml of puromycin for 4 hrs. Cells were stained for UBXN1 and ubiquitin. ALIS: Aggresome-like induced structures. (E) Aggresomes formed by Btz treatment contain primarily newly synthesized ubiquitylated proteins. Co-treatment with the translational inhibitor cycloheximide results in the loss of aggresomes in Btz treated cells. Cells were stained for ubiquitin and nuclei (Hoechst dye). (F) Immunoblot of global ubiquitin conjugate levels corresponding to samples in (E). Scale bar: 10μm.

**Supplementary Figure 3.**
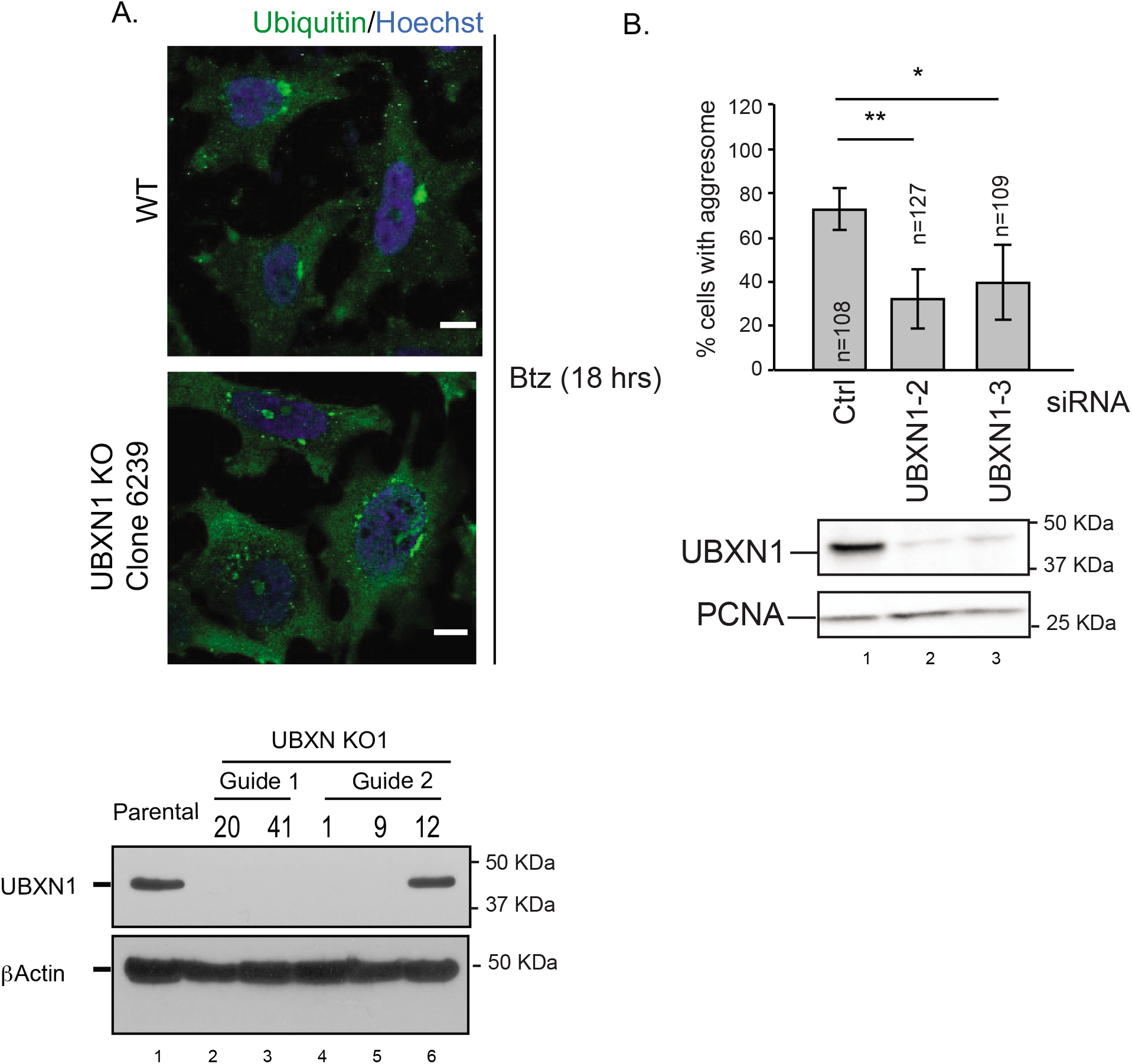
UBXN1 is required for the formation of aggresomes. (A) Parental or UBXN1 knock-out (KO) clone 6239 in HeLa Flp-in TRex cells were treated with 1 μM Btz for 18 hrs. Cells were stained for endogenous ubiquitin and nuclei. Bottom Panel: immunoblot showing loss of UBXN1 in CRISPR-Cas9 generated knockout clonal cell lines (Clone 9 is 6239 in panel A). (B) Quantification of aggresome formation upon transient depletion of UBXN1 with two separate siRNAs. Graphs show mean +/- standard deviation. Bottom Panel: immunoblot of transient depletion of UBXN1. The indicated number of cells was analyzed from the three independent biological replicates. *: p<=0.1, **: p<=0.05, as determined by unpaired students t-test. Scale bar: 10μm

**Supplementary Figure 4.**
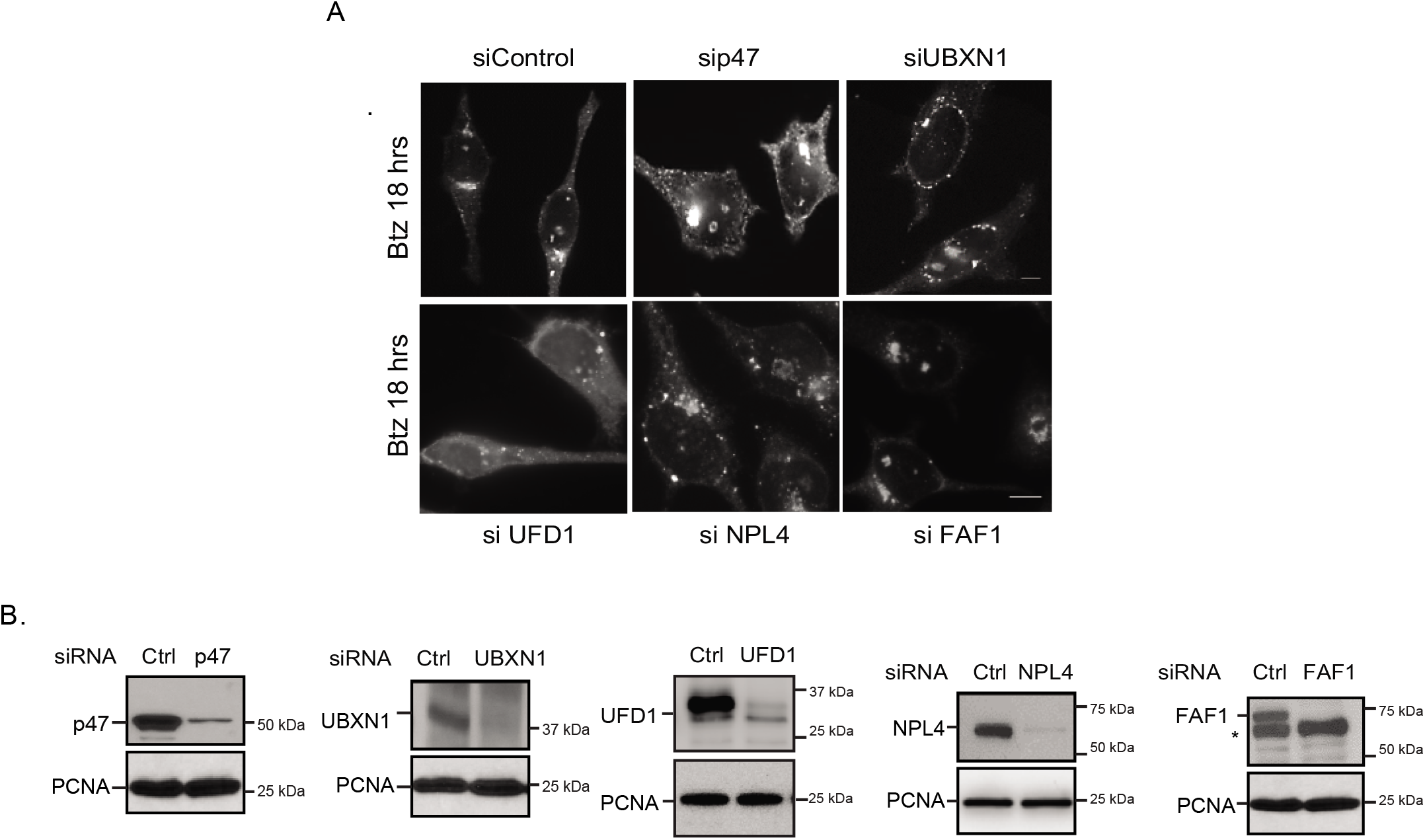
Specificity of aggresome formation phenotype. (A) Aggresome formation in Hela Flp-in T-Rex cells upon transient depletion of the indicated p97 adaptors. Cells were stained with anti-ubiquitin. (B) Levels of p97 adaptor depletion in Hela Flp-in TRex cell lines. Scale bar: 10μm.

**Supplementary Figure 5.**
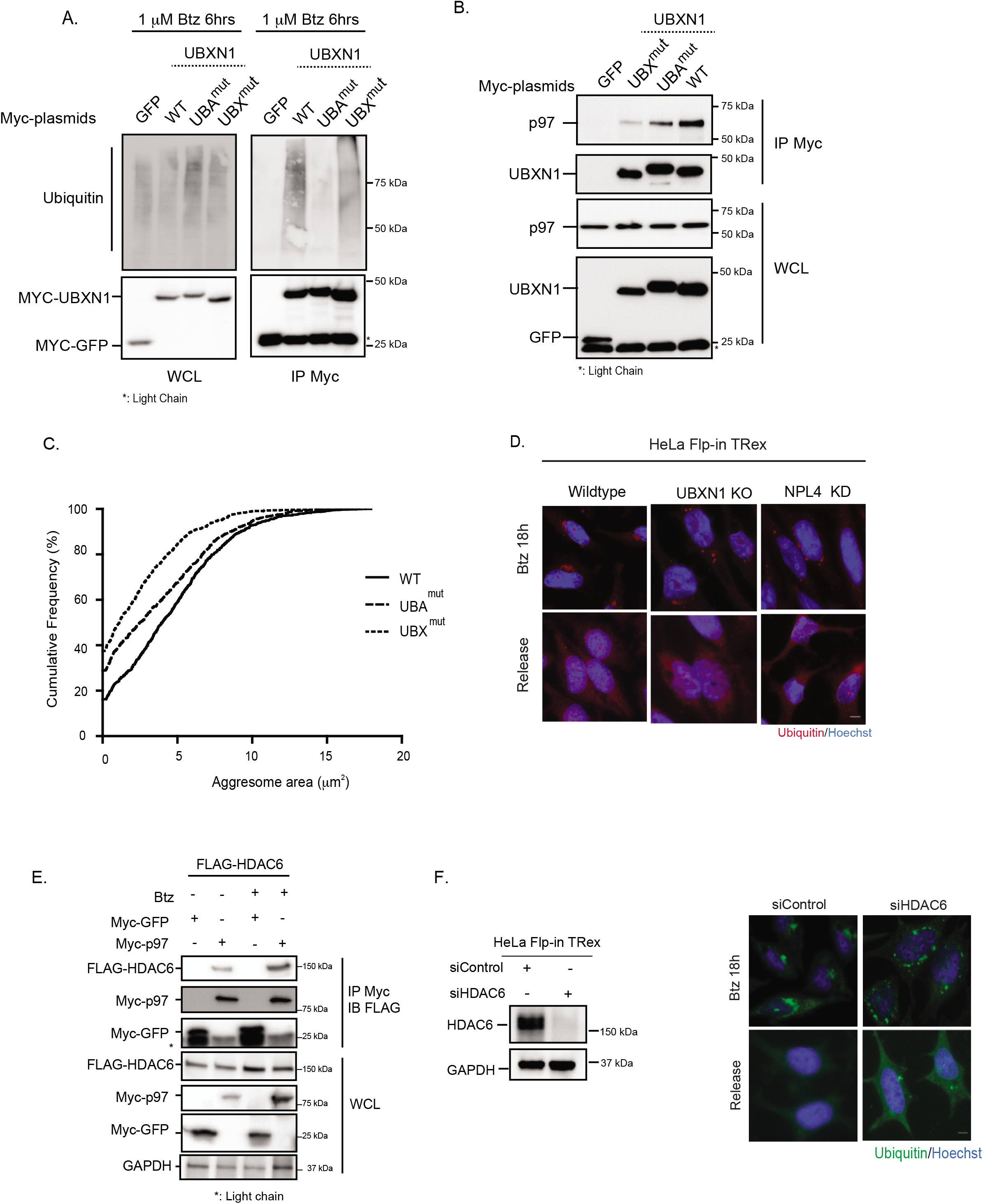
Characterization of UBA and UBX domains in UBXN1. (A and B) HEK-293T cells were transfected with the indicated Myc-tagged UBXN1 constructs. Cells were treated with Btz (A) and Myc affinity purifications were probed for endogenous ubiquitin (A) or p97 (B). (C) Cumulative frequency distribution of largest perinuclear aggregate areas in UBXN1 KO cells expressing GFP-UBXN1 wildtype, UBA, or UBX mutants. The left-ward shift of the curve indicates decrease in perinuclear aggregate area in the mutants (D) Wildtype, UBXN1 KO, or NPL4 KD cells were treated with Btz for 18 hrs and released into drug-free media to induce clearance of aggregates. The loss of UBXN1 or NPL4 did not impact aggregate clearance. (E) p97 interacts with HDAC6. HEK-293T cells were transfected with the indicated cDNAs and treated with Btz for 18 hrs, Myc affinity purifications were performed and probed for the indicated tagged proteins. (F) The depletion of HDAC6 inhibits aggresome formation and clearance in HeLa Flp-in TRex cells treated with Btz for 18 hrs. Scale bar: 10μm.

**Supplementary Figure 6.**
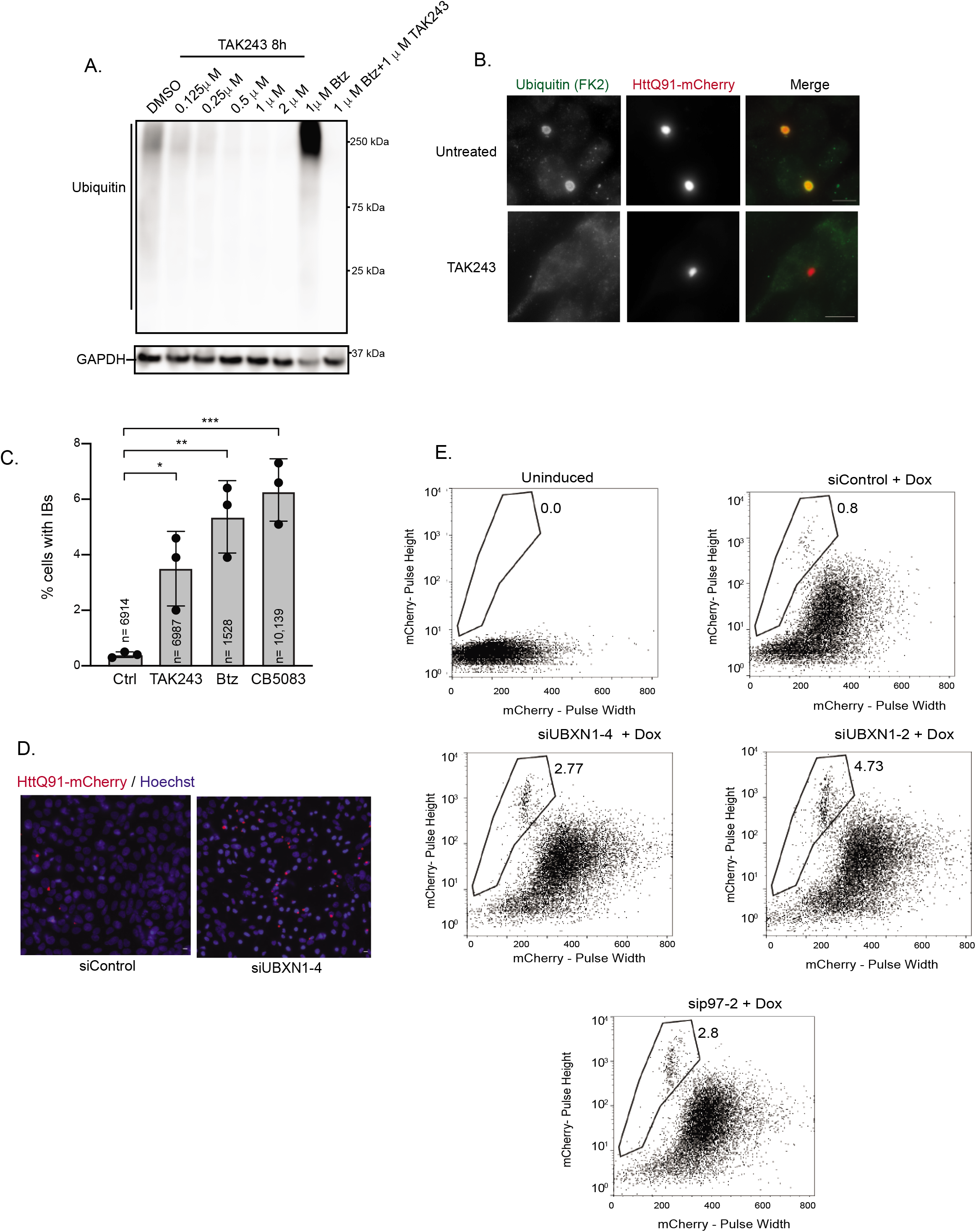
Role of p97-UBXN1 in HTT polyQ inclusion body formation. (A) Hela Flp-in TRex cells were treated with the indicated concentration of the ubiquitin E1 inhibitor TAK273 or Btz for 8 hours. Cell lysates were probed for endogenous ubiquitin. (B) U2OS cells were treated with doxycycline to induce the expression of HTT Q91-mCherry. Cells were treated with 1 μM TAK273 for 8 hours, fixed and stained for ubiquitin. Inclusion bodies still form but are devoid of ubiquitin staining. (C) Quantification of inclusion bodies observed by imaging in cells treated with 1 μM TAK274, 1 μM Btz, and or 10 μM CB-5083 for 8 hours. (D) Depletion of UBXN1 leads to an increase in HTT Q91-mCherry inclusion bodies. Quantification is provided in Figure 6B. (E) PuLSA analysis of HTT Q91-mCherry aggregates in UBXN1 and p97 depleted cells. The gates show the distribution of polyQ aggregates of various sizes based on pulse height. The greater the pulse height, the larger the inclusion body and vice versa. Note that this population represents only the HTT aggregates and not the complete mCherry signal. Total mCherry signal is represented in Figure 6C. The indicated number of cells was quantified in three biological replicates shown by the black dot. Graphs show the mean and standard deviation. *: p<=0.1, **: p<=0.05, ***: p<=0.001 as determined by One-way ANOVA with Bonferroni correction. Scale bar: 10μm.

